# An interpretable connectivity-based decoding model for classification of chronic marijuana use

**DOI:** 10.1101/2021.05.04.442433

**Authors:** Kaustubh R. Kulkarni, Matthew Schafer, Laura Berner, Vincenzo G. Fiore, Matt Heflin, Kent Hutchison, Vince Calhoun, Francesca Filbey, Gaurav Pandey, Daniela Schiller, Xiaosi Gu

**Author notes:** Authors contributed equally (Kaustubh R. Kulkarni, Matthew Schafer). **Corresponding author:** Xiaosi Gu, Center for Computational Psychiatry, Icahn School of Medicine at Mount Sinai, 55 W 125^th^ St, New York, NY 10027.

## Abstract

**Background:** Psychiatric neuroimaging typically proceeds with one of two approaches: encoding models, which aim to model neural mechanisms, or decoding models, which aim to predict behavioral or clinical characteristics from brain imaging data. In this study, we seek to combine these aims by developing interpretable decoding models that offer both accurate prediction and novel neural insights. We demonstrate the effectiveness of this combined approach in a case study of chronic marijuana use.

**Methods:** Chronic marijuana (MJ) users (n=195) and non-using healthy controls (n=128) completed a cue-elicited craving task during functional magnetic resonance imaging. Linear machine learning methods were used to classify individuals into chronic MJ users and non-users based on task-evoked, whole-brain functional connectivity. We then used graph theoretic analyses to identify ‘predictive functional connectivities’ among brain regions that contributed most substantially to the classification of chronic marijuana use.

**Results:** We obtained high (~80% out-of-sample) accuracy across four different classification models, demonstrating that task-evoked, whole-brain functional connectivity can successfully differentiate chronic marijuana users from non-users. Subsequent network analyses revealed key predictive regions (e.g., anterior cingulate cortex, dorsolateral prefrontal cortex, and precuneus) that are often implicated in neuroimaging studies of substance use disorders, as well as some key exceptions. We also identified a core set of networks of brain regions that contributed to successful classification, comprised of many of the same predictive regions.

**Conclusions:** Our dual aims of accurate prediction and interpretability were successful, producing a predictive model that also provides interpretability at the neural level. This novel approach may complement other predictive-exploratory approaches for a more complete understanding of neural mechanisms in drug use and other neuropsychiatric disorders.

## INTRODUCTION

Psychiatric neuroimaging has two main goals: describing the neural mechanisms of mental dysfunction and predicting clinical characteristics from neural data^1^. These goals are typically approached with different statistical and inferential paradigms, each with their own different strengths and weaknesses^2^. Common functional magnetic resonance imaging (fMRI) modeling approaches allow investigations into the neural mechanisms of psychiatric disorders by testing hypotheses about how mental processes are represented in brain signals. Such “encoding” approaches model brain activity as a function of different features [i.e., estimating *p*(*Brain Activity*|*Features*)., or probability of brain activity conditioned upon features], but do not easily yield inferences about processes or clinical categories from brain activity [i.e., *p*(*Features*|*Brain Activity*)]. Given the functional diversity of the brain regions implicated in psychiatric disorders, establishing the functional specificity of a brain signal is difficult and limits the ability of encoding models to predict clinical characteristics from brain imaging data^3^.

In contrast, “decoding” models provide the opposite type of inference, as in these models, neural data are used to predict features^2^, such as clinical diagnosis [i.e., *p*(*Diagnosis*|*Brain Activity*)]. Machine learning (ML) models are often used for this purpose; in psychiatry, and substance use disorders specifically, numerous machine learning approaches have been used, including support vector machines^4–7^, logistic regression^8–10^, and others^11–13^. However, decoding models do not necessarily give insight into neural mechanisms, or even neurobiological plausibility^2^ and are generally considered less interpretable than encoding models.

In recent years, there have been numerous attempts to unify the descriptive and predictive approaches: by linking patterns of brain activity to known variations between perceptual task stimuli^14–16^, by aligning highdimensional functional brain data and subsequently decoding in the aligned space^17–20^, and many others^2,21–23^. Although each of these classes of models allows unique insights into brain functions, they are still fundamentally different from the approach we propose here, which explicitly generates a predictive model first, and only applies interpretation analyses to the model weights. This serial approach of training and subsequent interpretation is known to be a challenging, understudied, and highly important goal in machine learning^24–26^, especially in the field of neuroimaging research^27,28^.

One way to improve decoding models’ interpretability is through theory-based modeling decisions about the types of neural features on which to train the model (feature selection)^27,29,30^. For example, there may be more information about psychiatric dysfunction in the interactions between regions than in the activities of isolated regions^31^. These competing hypotheses can be tested in the same data (model comparison): models trained features with more relevant information should produce better predictions.

Another way to gain insight from decoding models is to analyze the model weights. Decoding models trained on functional connectivity indicate the features of network activity that are predictive of the outcome. One novel approach is to apply network analysis to understand the predictive (i.e., weighted) connectivity. In recent years, network neuroscience has emerged as a powerful tool to provide essential metrics and methods to uncover complex brain interactions^32–35^. We employ these network analytic methods to infer brain structures critical for accurate classification. Importantly, the inferences we draw about group differences in network features are constrained by the predictive performance of the decoding model [i.e., *p*(*Network Features*|(*Diagnosis*|*Connectivity*))].

In this study, we use a large fMRI dataset^36,37^ collected from individuals with and without chronic MJ use (i.e., cannabis use disorder). To our knowledge, this is, to date, the largest fMRI sample used in the classification of substance use disorders (n=323), the first attempt to classify chronic MJ use with fMRI, and the first utilization of network analysis to interpret a fMRI decoding model. In recent years, reduced perception of adverse effects of cannabis has coincided with increased usage and legalization efforts^38–41^. Although the adverse clinical effects of cannabis have been well-established^39,42–45^, research on them has been hampered by the absence of reliable mechanistic biomarkers of cannabis use disorder. With our predictive and interpretable modeling approach, we aim to address this critical gap in knowledge.

We present here a novel modeling approach to balance the dual goals of clinical prediction and mechanistic understanding. We trained linear decoding models on whole brain functional connectivity from individuals with chronic use and healthy controls during a marijuana cue-induced craving paradigm. The models predicted chronic use of MJ with high accuracy in out-of-sample participants (~80%) and outperformed models that used only regional activities - suggesting that the interactions between brain regions contained more information about the differences between these groups. Network analyses on the predictive connectivity matrices (i.e., functional connectivity weighted by the model coefficients from predictive models) identified brain regions and networks important to successful use classification, demonstrating the utility of interpretable decoding models for neurobiological description.

## RESULTS

### Model training: classification of chronic marijuana use

We first trained decoding models using two different linear machine learning algorithms (logistic regression [LR] and linear support vector machine [SVM]) to predict the clinical label of chronic marijuana use from wholebrain functional connectivity. Two regularization penalty types (L_1_ and L_2_ penalty) were chosen to be applied to each learning algorithm, for a total of four candidate classification algorithms. The algorithms’ inputs consisted of a 4,005-element vector representing pairwise correlation values between every region in the brain as defined by the Stanford 90 region of interest (ROI) atlas (see **Methods** for more details). The full dataset (two runs each from n=195 chronic marijuana users, n=128 non-users) consisted of 646 total runs. Subjects were divided into training and testing splits: 80% for training (258 subjects, 516 samples), and 20% (65 subjects, 130 samples) for out-of-sample testing. The 80% training set was used to optimize our chosen hyperparameter (regularization penalty strength [α]) across the algorithms and penalty types, with 10-fold cross-validation. The 20% testing set was used validate the performance of the best-performing models from training. The complete computational analysis pipeline is depicted in **Fig. 1**. Cross-validated accuracies for each combination of hyperparameters are summarized in **Table 1**.

**Fig 1.**
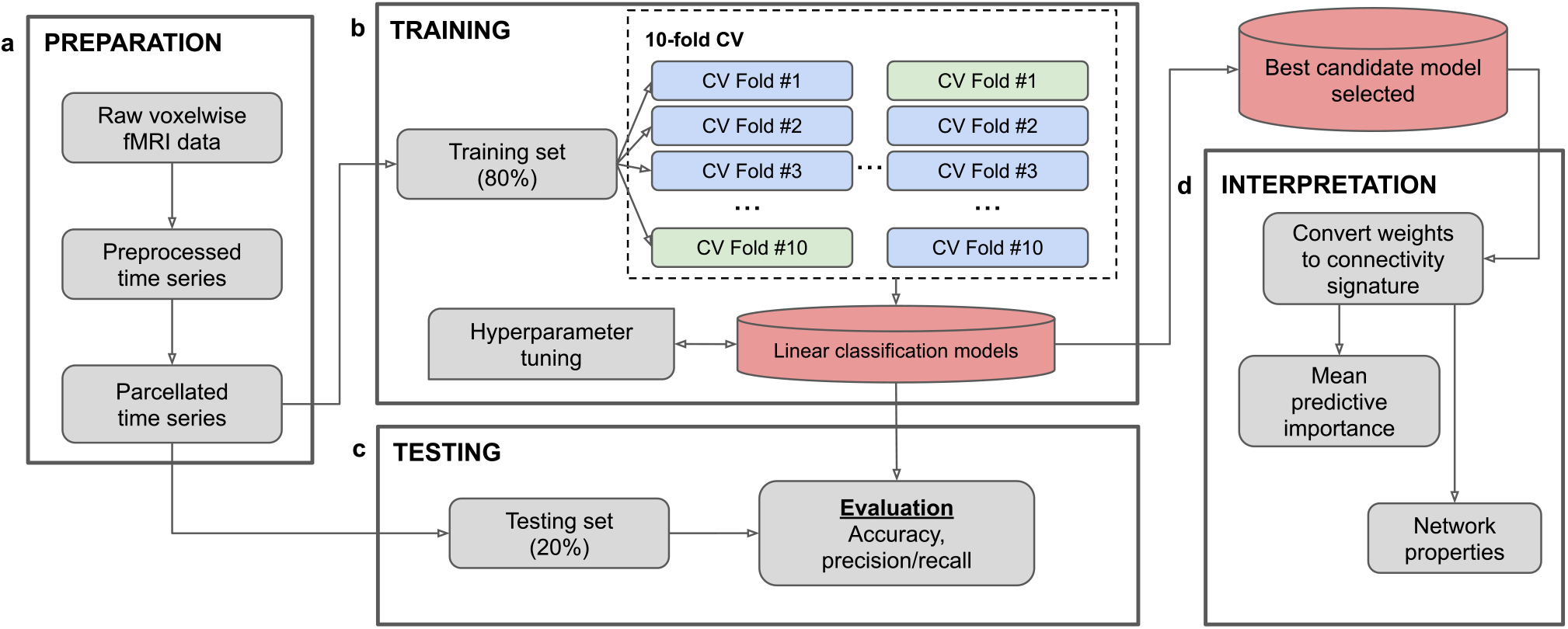
Machine learning pipeline. **(a)** Raw voxelwise time series are preprocessed using the fmriprep preprocessing pipeline. Minimally preprocessed files are brain-masked and smoothed with a 4mm FWHM Gaussian kernel. Nuisance/task regression is performed (see Methods for list of regressors used). Clean voxelwise time series is parcellated into 90 functional ROIs using Stanford functional atlas. **(b)** Parcellated data are divided into 2 sets; the training set is used for training and cross-validation, the testing set is used to evaluate the optimized classification models (shown in the cylinders). The optimization set is further divided into 10 subsets for cross-validation. Four linear classification algorithms are selected for hyperparameter tuning (L1, L2 penalized logistic regression and linear support vector classification). An alpha hyperparameter, corresponding to regularization strength is selected cross-validated accuracy as a metric. **(c)** The optimized hyperparameter tuned model is re-trained with the full training dataset and evaluated using the testing dataset. Evaluation parameters include accuracy, and precision/recall scores. (d) The best performing model (shown in the cylinder) is then trained on the full dataset (training + testing) to prepare for interpretation analysis. The weights derived from the linear models are converted to a connectivity signature and used to characterize brain connectivity structures important for prediction of chronic cannabis use. This analysis includes a regional mean predictive importance metric, as well as network characterization of subject-specific connectivity matrices weighted by the model weights.

**Table 1.** Hyperparameter optimization: prediction accuracy. Algorithm prediction accuracy is compared with 10-fold cross-validation of the training set (516 subjects) varying two hyperparameter domains: penalty type and regularization strength. Higher alpha values correspond to higher regularization. Results show that low regularization strength works most effectively across all penalty types. Generally, L_1_ and L_2_ penalties work equally well at low regularization and L_2_ outperforms L_1_ at high regularization. A regularization value of α=0.0001 for both classification methods and penalty types were chosen for subsequent analyses.

| Regularization strength ($\alpha$) | L<sub>1</sub> LR | L<sub>2</sub> LR | L<sub>1</sub> SVM | L<sub>2</sub> SVM |
| --- | --- | --- | --- | --- |
| 1e-10 | 0.727 | 0.734 | 0.754 | 0.754 |
| 1e-7 | 0.733 | 0.742 | 0.740 | 0.742 |
| 1e-4 | 0.761 | 0.722 | 0.750 | 0.740 |
| 0.1 | 0.604 | 0.746 | 0.604 | 0.759 |
| 1 | 0.605 | 0.694 | 0.605 | 0.680 |
| 10 | 0.482 | 0.612 | 0.482 | 0.609 |
| 100 | 0.482 | 0.601 | 0.482 | 0.618 |
| 1000 | 0.482 | 0.482 | 0.482 | 0.482 |

### Model training: hyperparameter performance

Generally, the L_2_ penalty yielded with better performance for both LR and SVM, and lower α (corresponding to lower regularization strength) improved model performance, indicating that widespread information from many pairwise region correlations contributed to classification success. Based on these results, we selected 0.0001 as the value of α for the following analyses, given its reliably strong cross-validated performance across all penalty types and classification algorithms. Since both L_1_ and L_2_ penalties for both algorithms performed well for a range of α values, we used both for the final evaluation of the LR and SVM models on the training and testing sets (2 models x 2 penalties x 1 α level = 4 tested models). The receiver operating characteristic (ROC) curves shown in **Fig. 2** demonstrate the classification performance of these four models across various decision thresholds within the training set.

**Fig 2.**
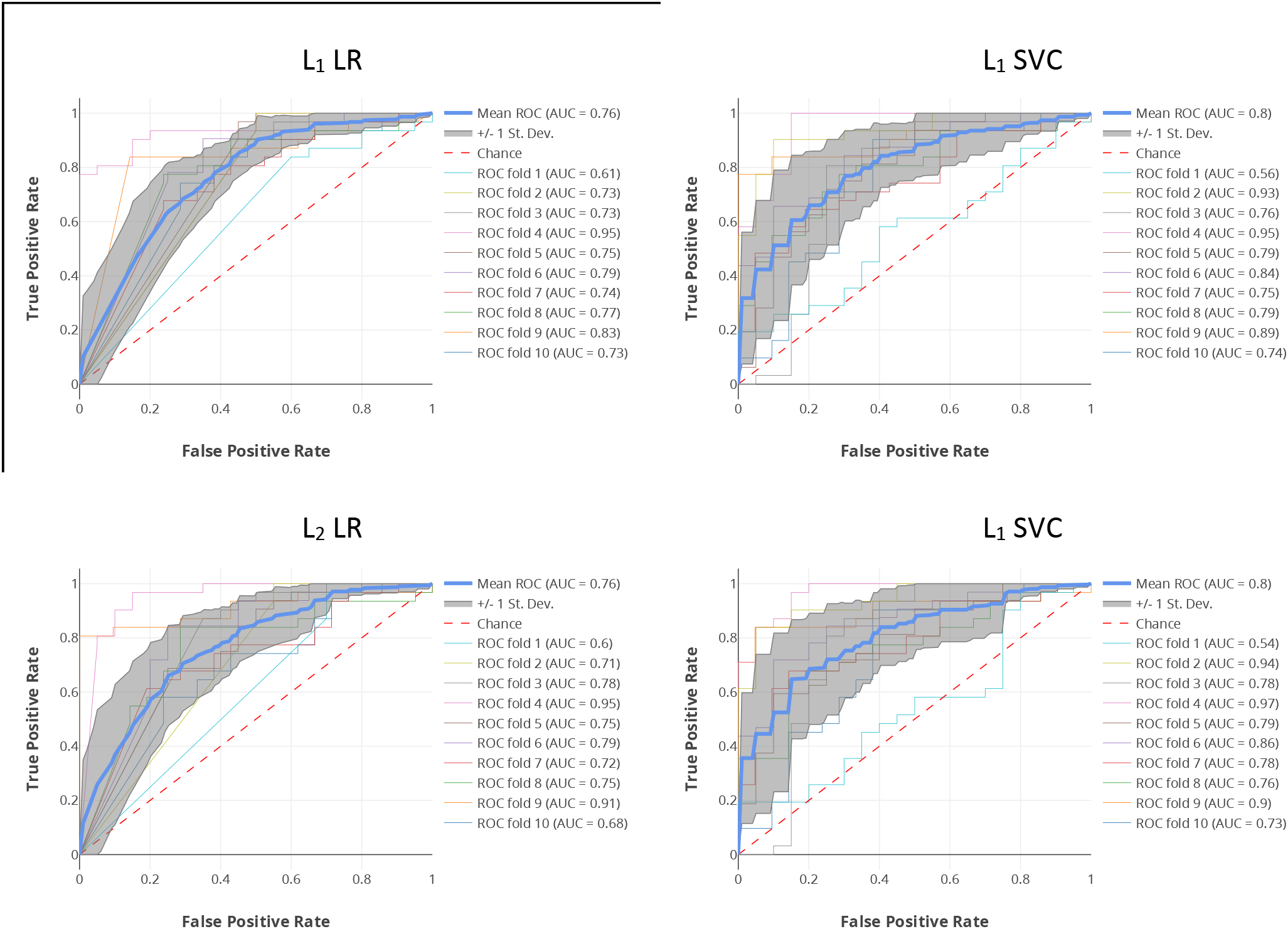
Receiver operating characteristic (ROC) curves for linear classification algorithms. The performance in 10-fold cross-validation shown for the four models at α=0.0001 was examined. All four models performed well in cross-validation metrics with the mean receiver operating characteristic curve well above chance (red dotted line). The logistic regression algorithm returns class probabilities that can be directly mapped to the ROC. The linear support vector classification algorithm returns only a decision function, corresponding to the signed distances to the hyperplane. These distances are converted to probabilities using Platt’s method.

### Model training: connectivity-vs. activity-based models’ performance

To test our hypothesis that models trained on functional connectivity would have higher prediction accuracy than models trained on regional activities, we repeated the same model training and evaluation process as described above with models trained on mean regional activation distances – the pairwise absolute value differences between regions’ mean estimated activity – as model inputs. Mean distances between the estimates were used rather than the estimates themselves to keep the number of features consistent across the models. Three different types of regional activity estimates were tested: mean time courses, mean marijuana cue betas (mj_cue_) and the contrast of mean marijuana cue betas minus mean control cue betas (mj_cue_>ctl_cue_). In the cross-validation on the training set, the highest performing models from each type of estimate (time course: 60.4%, mj_cue_: 65.1%, mj_cue_>ctl_cue_: 60.5%) had substantially lower accuracy than the best functional connectivity models. This result supported our hypothesis that functional correlations are more informative than isolated activities for differentiating chronic MJ users from healthy controls.

### Model testing: performance on held-out data

Next, we evaluated the out-of-sample performance of the final four models on the previously held-out test data (**Fig. 1c & d**). The resultant performance metrics are summarized in **Table 2** in terms of accuracy, AUC, and precision and recall for each model. Note that the accuracies for these models are much higher than chance testing set accuracies obtained by simpler/randomized models of three types: models that simply predict the dominant class (60%) or the averages of the accuracies of the random models generated after shuffling the training subjects’ labels 1000 times (~53% for all the models).

**Table 2.** Out-of-sample performance metrics. Out-of-sample (OOS) performance metrics are summarized after re-training each model on the full training set (379 subjects). The precision and recall values reported here are the average of both classes, weighted by the number of participants in each class.

|  | **L<sub>1</sub> LR** | **L<sub>2</sub> LR** | **L<sub>1</sub> SVM** | **L<sub>2</sub> SVM** |
| --- | --- | --- | --- | --- |
| **Accuracy** | 0.777 | 0.754 | 0.792 | 0.777 |
| **Area Under Curve** | 0.772 | 0.728 | 0.779 | 0.756 |
| **Precision** | 0.780 | 0.752 | 0.791 | 0.775 |
| **Recall** | 0.777 | 0.754 | 0.792 | 0.777 |

**Table 3.** Community membership. For each community identified, the brain regions within that community are listed along with their mean degree centrality (DC) across participants. Communities are o rdered by highest average DC within-community.

| **Community 1** | **Mean DC** |
| --- | --- |
| Bilateral medial posterior precuneus | 0.0635 |
| R middle frontal gyrus | 0.0591 |
| R mid-temporal cortex | 0.0511 |
| R inferior parietal/angular gyrus | 0.0473 |
| R inferior temporal cortex | 0.0472 |
| L middle thalamus | 0.0401 |
| R precuneus | 0.0398 |
| L inferior temporal gyrus | 0.0372 |
| L superior temporal gyrus | 0.0357 |
| R pre/post-central gyri | 0.0289 |
| L mid occipital cortex | 0.0256 |
| R inferior cerebellum | 0.0240 |
| L inferior parietal cortex | 0.0202 |
| R mid occipital cortex | 0.0198 |
| L cerebellum | 0.0194 |
| Bilateral medial precuneus | 0.0175 |
| R cerebellum | 0.0131 |
| **Community 2** |  |
| L pre/post-central gyri | 0.0504 |
| L superior temporal/auditory | 0.0411 |
| R superior temporal/auditory | 0.0310 |
| L crus cerebellum | 0.0239 |
| Cerebellar vermis | 0.0234 |
| Bilateral supplementary motor area | 0.0197 |
| L anterior cerebellum | 0.0109 |
| Bilateral calcarine cortex | 0.0108 |
| **Community 3** |  |
| R lateral angular gyrus | 0.0422 |
| Bilateral posterior cingulate | 0.0337 |
| Bilateral mid-posterior cingulate | 0.0288 |
| L medial angular gyrus | 0.0286 |
| R superior frontal gyrus | 0.0270 |
| L lateral angular gyrus | 0.0250 |
| L ventral precuneus | 0.0156 |
| R inferior frontal gyrus | 0.0096 |
| **Community 4** |  |
| Bilateral ACC | 0.0548 |
| Bilateral mPFC | 0.0379 |
| L middle frontal/dIPFC | 0.0321 |
| R middle frontal/dIPFC | 0.0274 |
| L inferior parietal/angular gyrus | 0.0211 |
| L fusiform gyrus | 0.0193 |
| L inferior frontal gyrus | 0.0174 |
| L superior frontal gyrus | 0.0167 |
| R frontal gyrus | 0.0158 |
| R middle orbito-frontal cortex | 0.0155 |

In addition to their high predictive accuracy, the models also produced very similar predictions, with high similarity for the pairwise comparisons (Jaccard similarity coefficients: 0.81-0.91), suggesting that all chosen algorithms converged on similar results. Together, these findings demonstrated that linear modeling of whole brain functional correlations was effective in classifying chronic marijuana use.

### Model interpretation: predictive connectivity

Next, we interpreted the classification models’ weights to infer the brain connectivity structure implicated in chronic marijuana use (see **Fig. 1d** and **Methods**). The learned model weights had high rank similarity (Kendall’s tau coefficients: 0.70-0.75), suggesting similar relationships were learned by the different algorithms. One advantage of using linear models is that the input features and learned model weights share the same shape: specifically, in our case, the model weights were the signed coefficients of region-to-region connectivity values that, when added, produced the classification decisions. This enables meaningful interpretability of the brain connectivity patterns most significantly implicated in differentiating chronic MJ cases from controls. First, we organized the model weights for each of our four linear models into a 90×90 pairwise region feature weight matrix (see **Methods**). Then we combined the model weight matrices with the pairwise correlation magnitudes, subject by subject, to produce weighted connectivity matrices - which we refer to as “predictive connectivity” (see **Fig. 3**). Finally, we used two model interpretation approaches: (1) We evaluated the regions with the highest predictive importance for each model, and (2) performed a graph theoretic analysis on the model weight matrices to examine network properties.

**Fig. 3.**
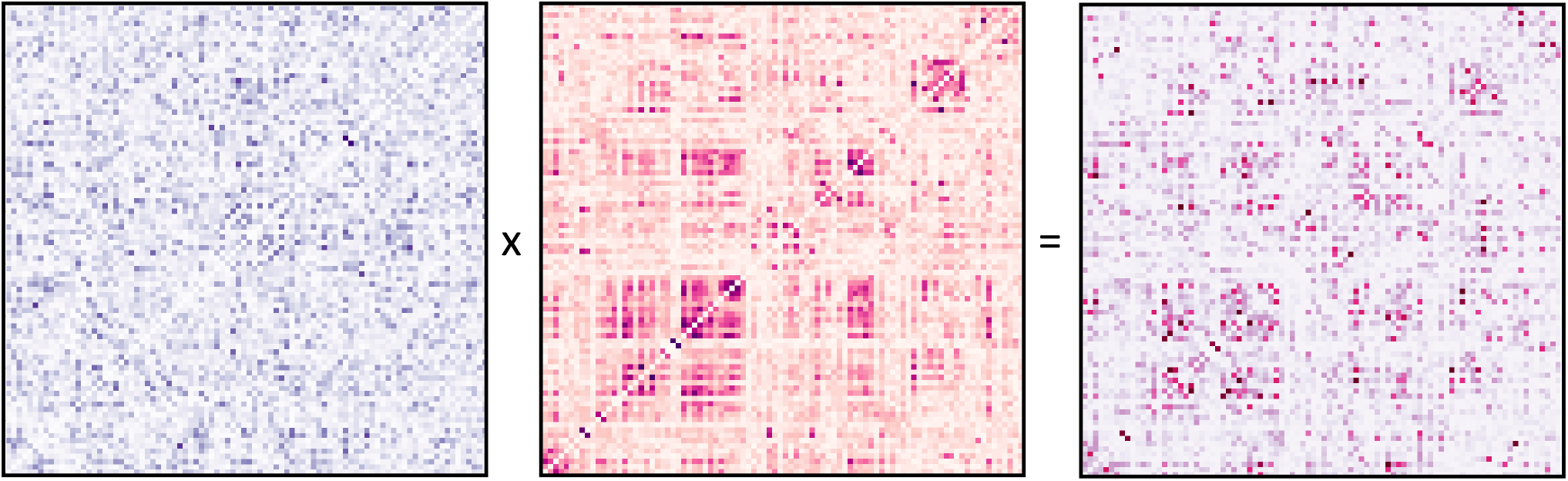
Generating subject-specific weighted connectivity matrices. For each participant, the absolute values of the functional connectivity were element-wise multiplied by the absolute values of the L_2_ SVM model weights to produce a weighted functional connectivity matrix (shown for a single, random subject). The weighted functional connectivity represents the importance of each model-weighted functional correlation to the resulting prediction for that participant: larger values represent a larger contribution.

#### Predictive importance of brain regions

First, we calculated the mean weighted correlation for each region with all other regions for every subject and ranked them by the average across subjects. Across the four models, we observed high consistency in the ranks of the regions in terms of predictive importance (Kendall’s tau coefficients=0.76-0.83, p-values=8.45e^-27^-4.71e^-31^). Given the similarities in weights, predictions, and regions of predictive importance across models, we assumed relative stability across the models and selected a representative for subsequent interpretations, namely the L_2_ SVM (α=0.0001) model due to its high and stable performance across the ranges of α tested. For this model, the top twenty regions in terms of mean predictive importance are shown in **Fig. 4a**. These include brain regions such as bilateral anterior cingulate cortex (ACC), left pre/postcentral gyri, right middle frontal gyrus, and right inferior parietal cortex.

**Fig. 4.**
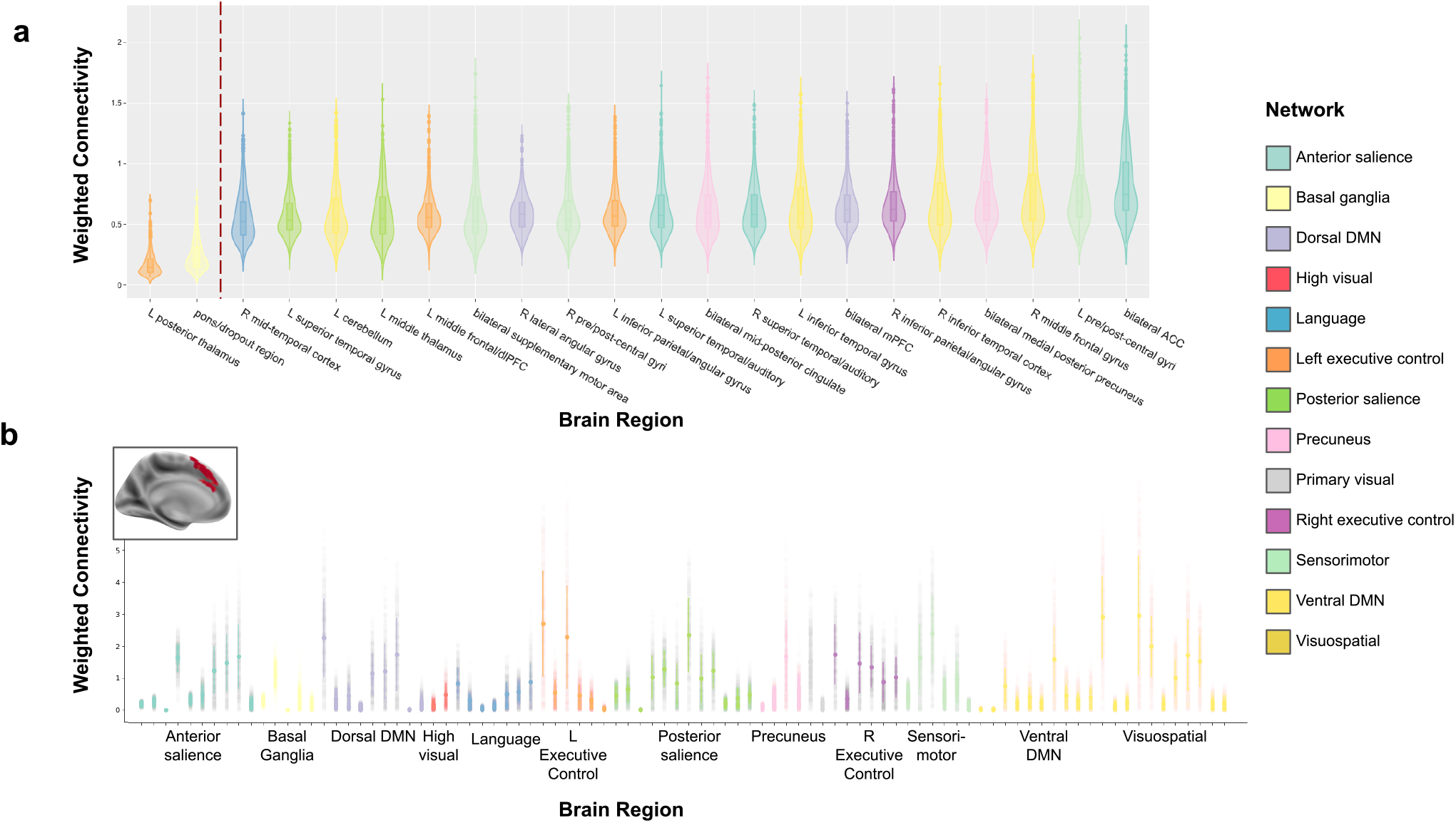
Top weighted averaged parcel connectivities for L_2_ SVM classifier. **(a)** Functional connectivity matrices were averaged across all subjects, and element-wise multiplication was performed with weights generated after model fitting with the L_2_ SVM algorithm. The resulting matrix was a weighted region-to-region connectivity matrix. The mean of absolute weighted connectivity was calculated for each region for each algorithm. The distributions (mean, quartiles and outliers) of the absolute weighted connectivities across all subjects is shown above. The top twenty regions with highest means of weighted absolute connectivity are shown on the right side of the graph, while the two lowest are shown on the left for comparison. In the axis labels, ‘L’ represents left lateralized, and ‘R’ represents right lateralized regions. Regions with the highest weighted connectivity include bilateral ACC, left sensorimotor cortex, middle frontal gyrus and bilateral angular gyrus. **(b)** For the regions identified as having high weighted connectivities, region-specific connectivity patterns were assessed at a group level. Here, the connectivity strength and direction are shown from bilateral ACC, the region with the highest weighted connectivity across participants, to every other region. ACC appears to have have high connectivity specifically to inferior, middle, and superior frontal cortical areas across multiple functional networks (executive control, ventral default mode, visuospatial) as well as precuneus/angular gyrus regions. This suggests the presence of an ACC + frontal cortex + lateral parietal cortex task network, later supported by our community detection analysis (see Fig. 10).

As an exemplar region, bilateral ACC showed high mean predictive importance across all models, so its unweighted regional connectivity strengths to every other region were further visualized in **Fig. 4b**. Among the regional connections to ACC, weighted connectivity correlated highly with original functional connectivity (r_mean_=0.478, r_std_=0.0628). Importantly, however, a number of regions showed relatively small magnitudes of connectivity strengths to ACC but high predictive importance (reflected by high model weights).

To examine whether top 20 regions identified by our top weighted connectivities (**Fig. 4a**) were consistent with those reliably implicated in craving, we compared our region-specific predictive importance scores i.e., ranked weighted connectivities, to uniformity and association maps retrieved from Neurosynth.org^46^ using a term-based meta-analysis. The ‘craving’ keyword yielded aggregated activation maps from 80 published studies thresholded at FDR-corrected p<0.01. The most significantly active regions identified by this meta-analytic approach included medial prefrontal cortex, middle cingulate cortex, medial prefrontal cortex, and medial parietal lobule. Each Neurosynth map was projected onto an anatomical map with the Stanford functional ROIs with high weighted connectivity (predictive importance) overlaid on top (**Fig. 5**). Map comparisons were restricted to a qualitative overview due to the highly dissimilar sparsity of the maps, as well as significant differences in the sizes of Stanford ROIs and the activation loci in the Neurosynth maps. We found that regions in the meta-analytic craving map qualitatively showed a moderate level of correspondence to regions identified as having high predictive importance, supporting our interpretation pipeline.

**Fig 5.**
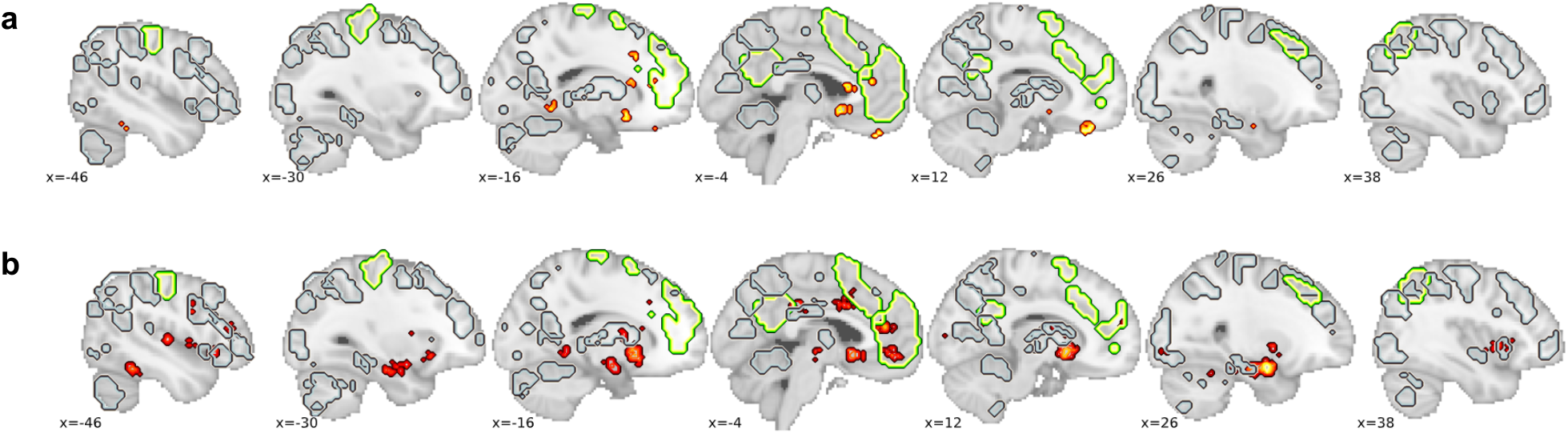
Meta-analytic comparison with Neurosynth craving maps. To compare our top regions of predictive importance to existing literature, we performed a direct comparison to association and uniformity maps retrieved from Neurosynth, a meta-analytic database. We used the ‘craving’ keyword to identify activations corresponding to all activations (uniformity) and unique activations (association) related to craving in the meta-analytic database. The average signal and proportion of voxels activated within-region was calculated and thresholded. Given the relative sparsity of the association map compared to the uniformity map, association was thresholded at 5% voxel participation and uniformity at 25% participation. These activations are shown in red, with (a) showing the association map and (b) showing the uniformity map. All the Stanford ROIs are overlaid on the map above at relevant sagittal slices (indicated with ‘x’), with green regions corresponding to ROIs identified as having high predictive importance in our analysis. There is a moderate level of overlap between the craving maps and our predictively important regions, demonstrating the utility of our approach in identifying regions grounded in previous literature, but also being able to generate new hypotheses for regions involved in distinguishing cannabis users from non-users.

#### Graph theoretic analysis

Our next goal was to investigate the networks of brain regions that were relevant to distinguishing chronic MJ users from non-users using a graph theoretic approach. Subject-specific graphs were generated by thresholding the weighted connectivity matrices described above (see **Fig. 3** and **Methods** for more details). Graph properties (see **Fig. 6**) were calculated for individual nodes in the graph (i.e., individual brain regions), the full graph (i.e., the entire whole-brain network), and clusters or sub-networks within the full graph that are highly modular (i.e., highly interconnected communities of brain regions).

**Fig 6.**
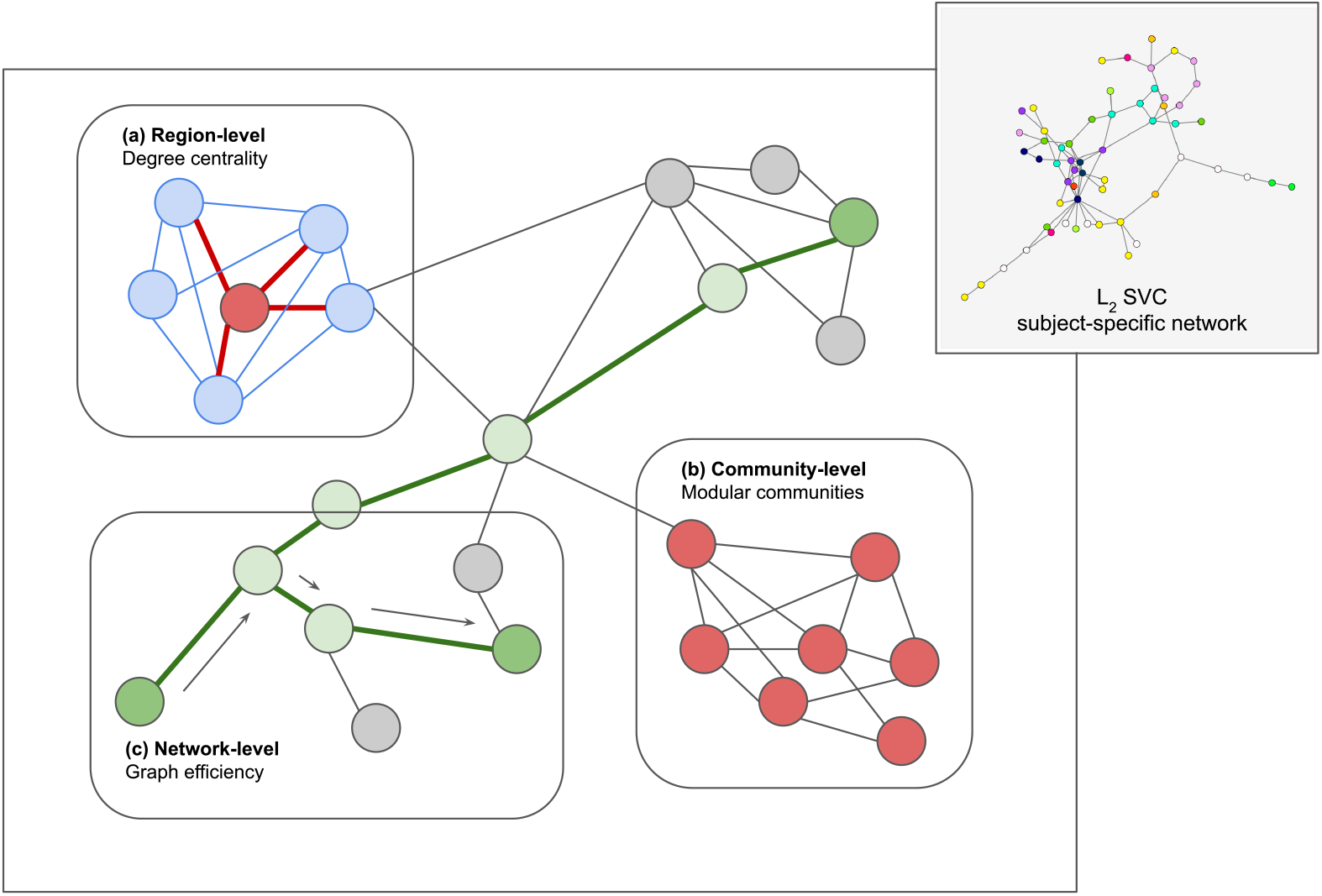
Network properties workflow. For each subject, a weighted connectivity matrix is generated by performing an element-wise multiplication of the original subject connectivity matrix and the model weights. The resulting matrices are thresholded to 2% sparsity to restrict to only highly informative connections. Network properties are then calculated at three different levels to characterize the subject-specific networks. (a) The degree centrality of each node of the network, i.e. a brain ROI, is obtained by calculating a normalized sum of surviving links to other nodes. In principle, this provides a measure of the importance of a region’s connections to other regions for prediction. (b) At the meso-level, community detection algorithms are used to divide the full network into modular sub-networks that are highly connected to each other. These communities correspond to brain patterns that together are highly important for prediction of chronic cannabis use. (c) At the network-level, global efficiency of the network is calculated by determining the inverse average shortest path. For each node, the distance to every other node is calculated and averaged. The process is repeated for every node and averaged across nodes. The inverse of this averaged shortest path length is the efficiency of the network. High efficiency networks exchange information well because they are densely connected, and thus have fairly low average path lengths.

At the brain (local) level, we calculated a brain region’s importance for prediction by calculating a subject-specific degree centrality (DC), or the number of predictively valuable connections a region has with other regions. Regions were ordered by their mean DC scores, indicating their levels of predictive importance. The top twenty regions of highest average DC (along with the two lowest for comparison) are shown in **Fig. 7**. Regions of high overall importance include bilateral ACC, right inferior parietal/angular gyrus, and right middle frontal gyrus. Regions from numerous resting state networks are represented in the top twenty regions, indicating widespread connectivity is important for distinguishing individuals with chronic MJ use from controls.

**Fig. 7.**
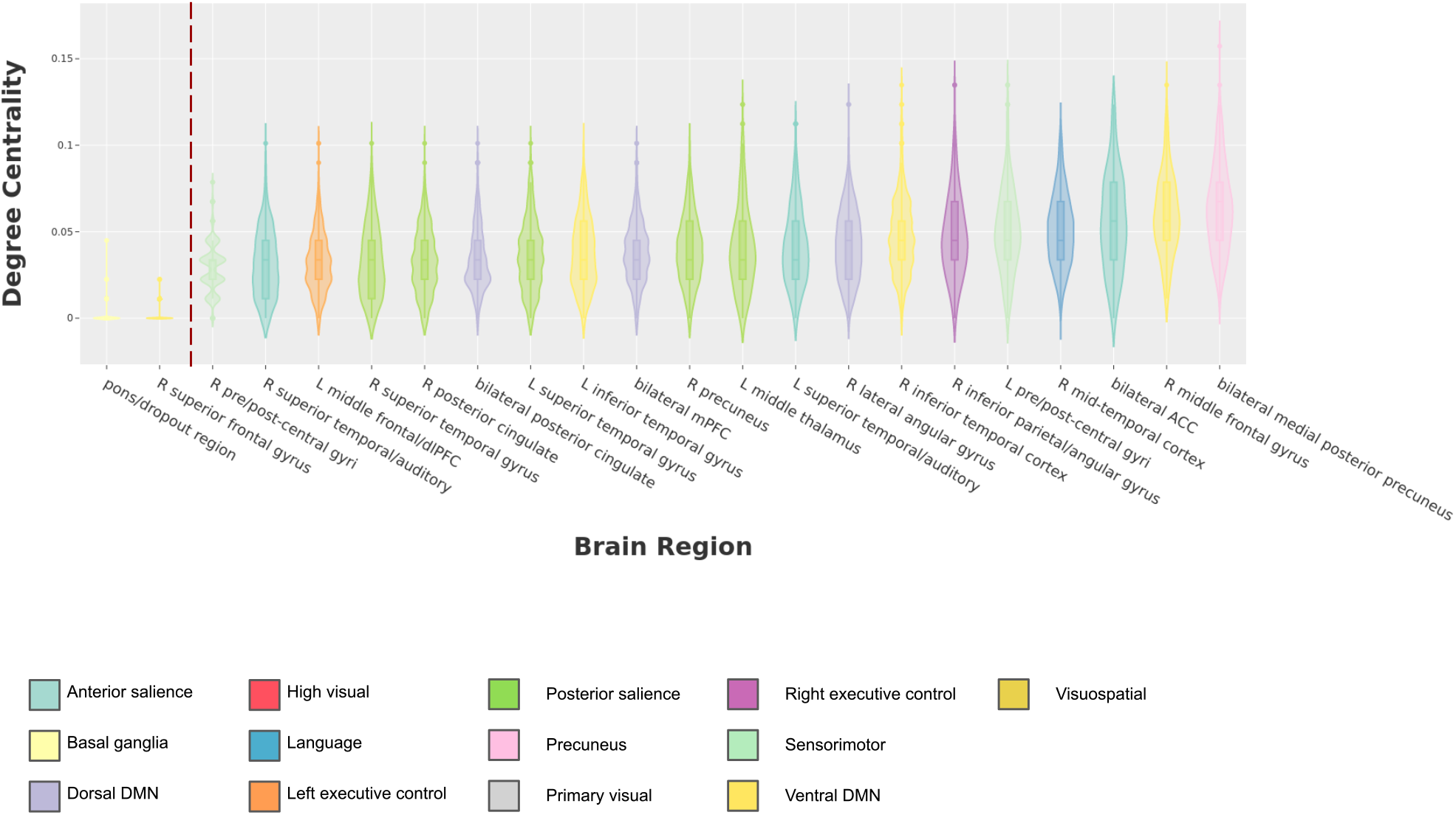
Subject-level degree centrality. Degree centrality represents the normalized number of weighted connections for each brain region that survive thresholding. In other words, it provides a measure of the level of distributed connectivity displayed by a brain region. In the plot above, degree centrality is calculated for each region independently, for each subject. The top twenty regions of highest mean degree centrality are shown, in addition to the lowest two for comparison. Regions identified as having high degree centrality across participants include middle frontal gyrus, bilateral ACC, and bilateral medial PFC. Note that there is a significant overlap here with regions identified as having highest absolute weighted connectivity (**Fig. 4**) but there are significant differences as well.

Next, we analyzed properties of network organization at the whole-brain (global) level by calculating network efficiency, or its ability to transmit information effectively. An independent samples t-test between the two groups was not significant, indicating that there were no differences in network efficiency for prediction of users vs. non-users.

Finally, at the community (meso) scale, we used community detection algorithms to discover modular subnetworks within the weighted connectivity matrices (see **Methods** for details). **Fig. 8a** shows the thresholded group-average weighted connectivity matrix reorganized by the discovered community structure. Each community was then ranked by its average degree centrality (DC) score. The highest ranked community included regions from bilateral ACC, bilateral supplementary motor area, right dorsolateral prefrontal cortex, and right inferior parietal/angular gyrus. The second top scoring community included right middle frontal gyrus, left angular gyrus, and bilateral medial precuneus regions. The top 4 modular communities that contributed most to prediction are visualized in **Fig. 8b**.

**Fig. 8.**
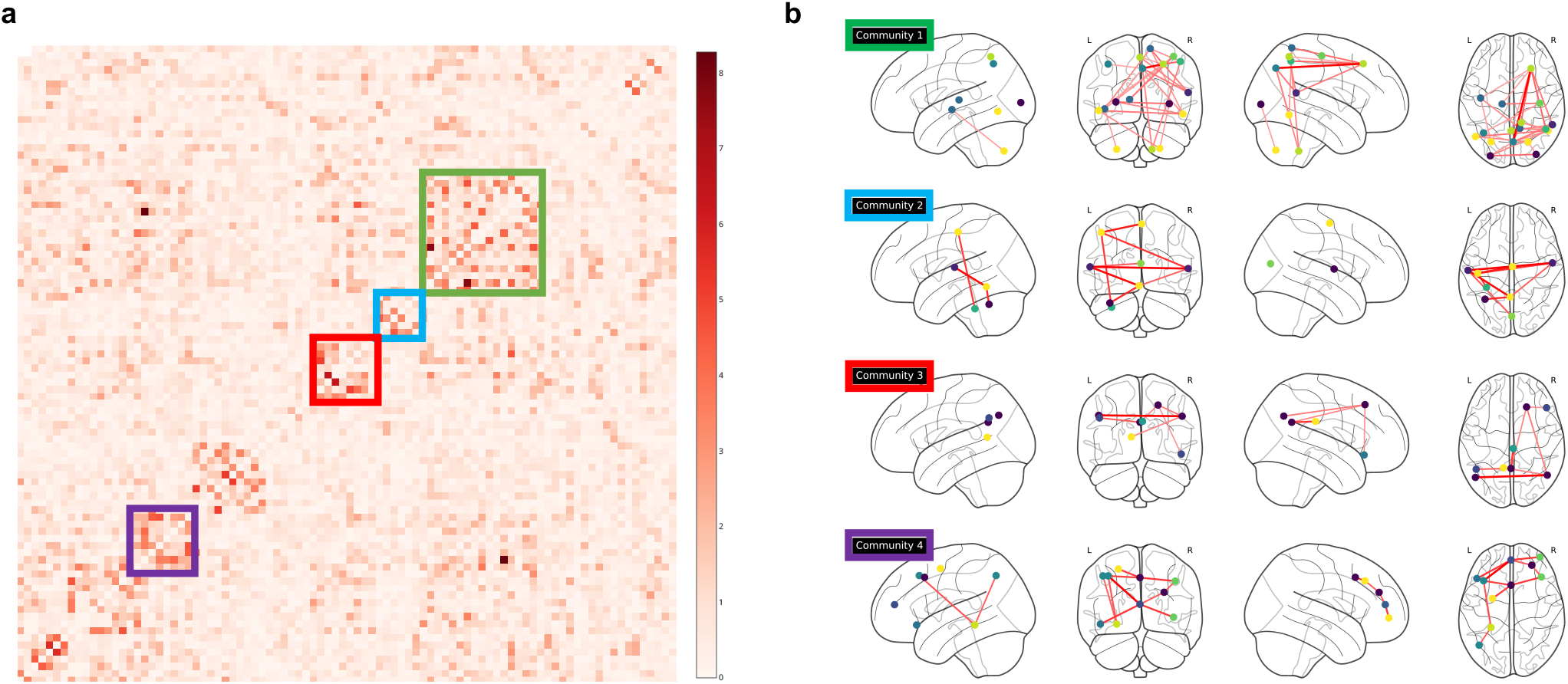
Predictive communities in group-average weighted connectivity matrix. The Girvan-Newman community detection algorithm was applied to the group-average weighted connectivity matrix. Girvan-Newman segregates communities within a group by iteratively removing edges with the highest betweenness centrality until a target modularity score is reached. Each disconnected set of nodes is then characterized as a community. **(a)** The group-averaged thresholded weighted connectivity is sorted by community assignment. Each colored square represents one of the top 4 communities by average degree centrality within community. **(b)** The color-corresponding communities are projected onto the brain and colored by resting-state network assignment as determined by the Stanford functional parcellation. The top 4 networks are largely bilateral. Community 1 is distributed mainly over posterior aspects of the brain and includes right pre/post-central gyrus, mid/superior temporal gyrus, precuneus, middle frontal gyrus and inferior parietal cortex. Community 2 contains cerebellar regions, superior temporal cortex, and left pre/post-central gyrus. Community 3 includes inferior and superior frontal gyri, angular gyri, and bilateral posterior cingulate. Finally, Community 4 includes bilateral medial PFC, bilateral dorsolateral PFC, bilateral anterior cingulate and right orbitofrontal cortex.

To confirm that this ranking reflected the predictive importance of each community, we performed a stepwise prediction analysis to determine the minimal number of communities necessary to produce good predictions. (**Fig. 9**) Starting with the regions in the highest DC ranked community, each region’s (non-redundant) pairwise correlations to all other brain regions were used to generate each participant’s distance to the hyperplane. With the inclusion of each additional community, the best performing decision threshold was determined in the training data and used to generate testing set predictions. The best testing set prediction came from the first 4 communities with 80% accuracy, outperforming even the overall model - and performing significantly better than random regions (permutation tested p=0.001).

**Fig. 9.**
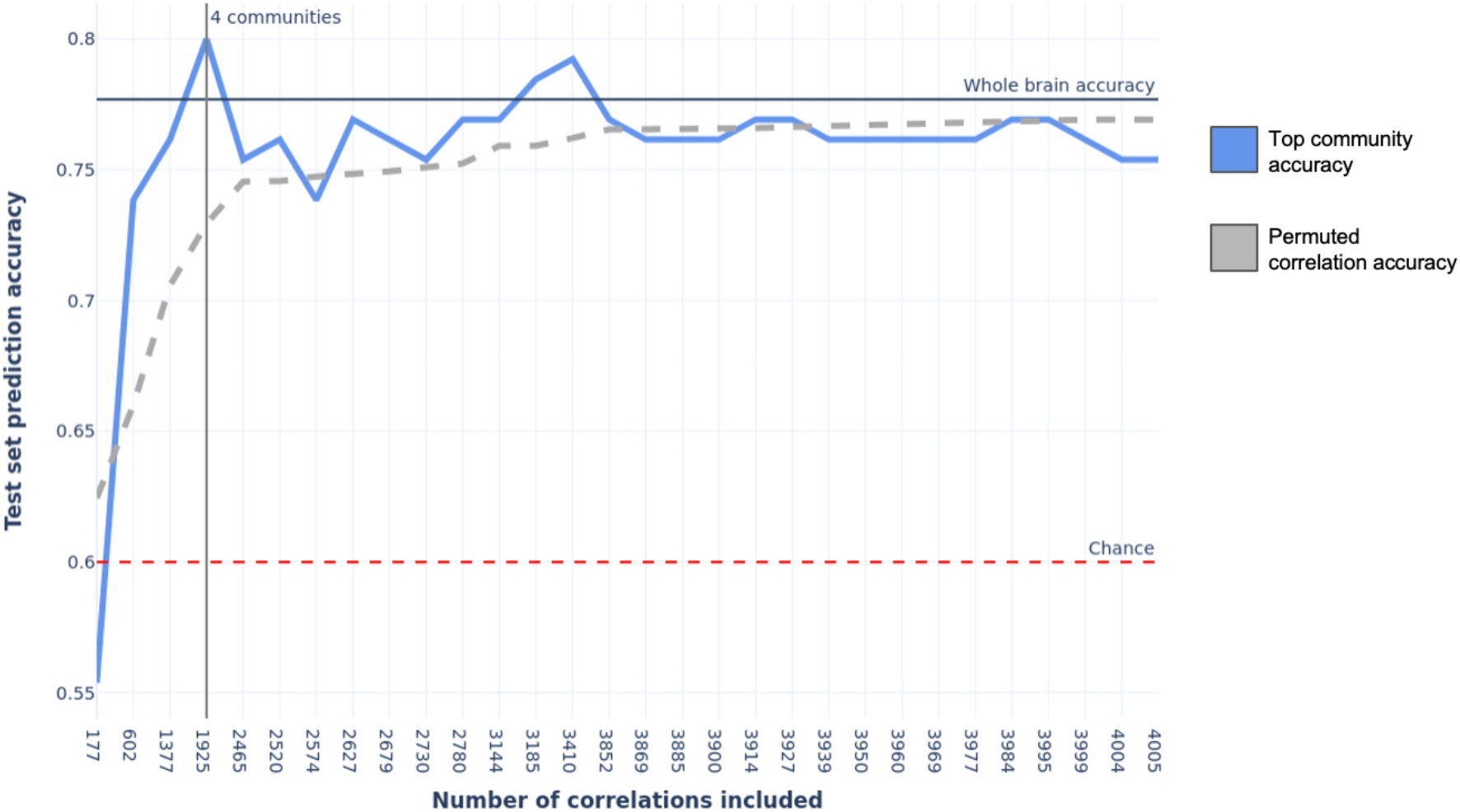
Predictive accuracies of top communities. A stepwise prediction analysis was performed to confirm the predictive importance of the top-ranked communities discovered in the community detection analysis. Starting with the highest ranked community, the correlations of all regions in the community to all other regions were used to generate distances to the hyperplane for each subject. Then, a search was performed for the optimal decision threshold that maximized prediction accuracy in the training data. Finally, this threshold was applied to the testing data to produce test set predictions. The best performing subset of communities was determined by the testing accuracy. Permutation testing (1000 permutations) was performed to judge the relative increase in performance using the top communities vs. using a random set of correlations while preserving the number of pairwise correlations included at each step. The permutation p-value was calculated as the percentile of the best performing non-permuted accuracy in the distribution of the 1000 permuted accuracies at that same step. Chance was defined as a naive classifier that always picks the dominant class (chance=0.60). The best testing set prediction came from the first 4 communities with 80% accuracy, performing significantly better than random regions (permutation tested p=0.001) and above chance.

## DISCUSSION

### Model testing: clinical prediction

In this study, we developed a novel modeling approach to balance accurate clinical prediction and model interpretability. Specifically, this approach classifies chronic marijuana users and healthy controls from task-based fMRI functional connectivity and subsequently identifies the individual regions and networks most important for this distinction. In the largest sample of individuals with long-term MJ use and healthy controls to date, we classified chronic use from functional connectivity during a cue-elicited craving task with nearly 80% out-of-sample accuracy. We used several different linear modeling approaches, all of which produced highly similar model weights, predictions, and regions with high mean predictive connectivity - suggesting they learned similar information. Our accuracies also compare favorably to previous fMRI decoding studies using functional connectivity to classify drug use, in both nicotine smoking^4,7,47^ and cocaine use disorder^48^ - even though most studies did not test out-of-sample or featured much smaller sample sizes (both of which can inflate prediction performance). Furthermore, this is one of the first fMRI study^49^, and the largest to date, to classify chronic MJ use (i.e., cannabis use disorder) - a relatively understudied drug use disorder.

### Model interpretation: predictive connectivity

Functional connectivity-based models outperformed models trained on regional activation estimates - suggesting there is more information about chronic MJ use in the interactions between regions than in their isolated activities. Given this, our next goal was to discover brain network patterns that differentiated the groups, starting with the individual regions that are most critical to successful prediction in the best performing model - the L_2_ linear SVC. Regions with high mean predictive connectivity were distributed across diverse resting state networks, such as the default mode, sensorimotor, salience and executive control networks - suggesting widespread functional differences between the healthy and MJ-using groups. Regions with widespread predictive connections were especially of interest and were judged by the number of functional connections between a region and the rest of the network that helped classify chronic use, so-called ‘predictive degree centrality’ (i.e., predictive DC). As shown in **Fig. 4b**, our interpretable model weights approach can even identify small magnitude connectivities between brain regions that are nevertheless highly important in differentiating chronic users from healthy controls.

There was high predictive DC in several sensory and motor related regions - including left inferior temporal gyrus, right inferior temporal cortex (both areas along the ventral visual pathway), bilateral primary somatosensory cortex and supplementary motor area. Given that the visual and tactile demands of the task were the same across groups, these regions likely reflect more than the passive reception of sensory information and output of motor commands. For example, these regions may facilitate the recognition of drug cues and retrieval of behavioral associations, such as the initiation of drug seeking/use behaviors^50^. Regions related to attention and its control also ranked highly on this measure - likely reflecting differential recruitment of attention during cue processing between the groups. For example, the right middle frontal gyrus, an important attentional control region and site of convergence for the dorsal and ventral attention networks^51^, had the highest predictive DC of any measured region. Bilateral ACC and dorsolateral prefrontal cortex (PFC), areas that feature dense cannabinoid receptors^52^ also ranked highly on this measure, corroborating previous reports of dysfunctional attentional and control-like processes during drug cue exposure and craving generally^53,54^ and in MJ users specifically^55,56^. High predictive DC was detected in regions associated with cue-reactivity and craving, including the precuneus and posterior cingulate cortex, regions that may work together to process drug cue salience and relevance to the self^57^ and in the bilateral medial PFC, which has extensive and recurrent dopaminergic connections with the ventral tegmental area and may direct drug-seeking behavior^58^. These findings suggest our method can recapitulate diverse findings from the literature.

We also discovered sets of brain regions (communities) that were important for to successful classification of chronic MJ users and controls and ranked them by average DC. The top four communities produced the best testing set prediction accuracy, even outperforming the inclusion of additional communities. These communities contained regions from different canonical resting-state networks (e.g., salience, default mode, frontoparietal networks), with two communities comprising the majority of predictively important regions. The first community included regions from bilateral ACC, posterior inferior temporal cortex and superior angular gyrus, and the second included regions from inferior angular gyrus, middle frontal gyrus, and superior temporal cortex. It is not obvious how these communities map onto canonical networks, suggesting that these potentially novel findings may reflect task-specific network organization.

The functional diversity of the regions and communities identified in these analyses suggests widespread functional differences between MJ users and controls - and the need for tasks that measure a wide range of structure-function hypotheses concurrently. It is possible that the relatively high accuracy we achieved in this study was due to the task: multiple sensory modalities and motor processes were engaged, allowing for more functional differentiation between individuals with MJ use and controls.

### Added value, limitations, and next steps

In general, decoding approaches use whole brain information during model fitting, culminating in a single statistical test, as compared to more standard encoding approaches (e.g., general linear modelling) that generally perform up to many thousands of tests across the brain and require extensive multiple comparisons correction. To the best of our knowledge, our proposed approach represents the first use of network analysis to interpret predictive models. Furthermore, our model interpretations are constrained on high decoding performance, conditioning our inferences upon the prediction of a real-world clinical label (self-reported behavior).

There are several limitations to this study. First, these classification accuracies reported are likely not high enough for direct clinical deployment. Further, the sample was divided only into chronic cannabis users versus non-users, not allowing us to disentangle continuous effects related to use. We also predicted a categorical label based upon self-report, not by real world behavior or underlying functional dimensions, and thus are bound by the accuracy of that label. This study also precludes most inferences about the specificity of the effects of marijuana use. Future work should compare marijuana users to chronic users of other drugs, as well as non-drug using individuals with other psychiatric dysfunction, in order to establish marijuana-specific neural signatures. Additionally, more data-driven parcellation approaches (e.g. using independent components analysis, gradient-based methods, or multimodal data) may elucidate more robust, replicable, or task-evoked neural signatures associated with chronic MJ use^59–61^. Another important direction for this approach clinically would be a longitudinal study predicting future risk of chronic use, especially in adolescents or young adults.

Many extensions to this joint predictive/explanatory approach are possible. The network analysis may be refined at the spatial scale, by generating voxel-level connectivity matrices and recalculating network properties. Another possibility would be to build predictive models from regions of interest, in a more hypothesis-driven manner (e.g., derived from areas of significant activation in an encoding model). Additionally, a regression-oriented predictive model would be an improvement over the classifiers outlined here: such approaches can make stronger inferences about the neural patterns of clinical features directly (e.g., symptom severity, craving), rather than indirect conclusions about patterns that differentiate clinical groups (i.e., chronic use or not)^2,21,27^.

This study is a first step towards building accurate and interpretable predictive models that have both theoretical and clinical significance. The models performed well in out-of-sample data, indicating their generalizability. Furthermore, we interpreted the best-performing model to both corroborate prior findings and discover potentially novel network-level properties in the context of substance use disorders. Future work can build on this approach of using joint predictive-explanatory models to constrain neurobiological inferences.

## MATERIALS AND METHODS

### fMRI data collection

#### Subjects

This study combined data from two pre-existing fMRI datasets (n=125 and n=198 respectively) measuring cue-elicited drug craving in participants recruited from the community (i.e. not treatment-seeking or inpatient) in Albuquerque, NM. The datasets include participants with and without chronic marijuana use (marijuana [MJ] n=195, healthy controls [HC] n=128 respectively)^36,37^. The combined data set has a mean age of 30, with 65% male participants.

#### fMRI scanner specifications

The two datasets had different fMRI scanner specifications, as described below.

**2009 sample**^36^: MRI images in this sample were collected in a 3T Siemens Trio scanner over two runs, for approximately 9 minutes and 22 seconds of scan time. T2* images were collected with a gradient echo, echo planar imaging protocol, with the following specifications: time to repetition (TR) of 2,000ms, time to echo (TE) of 27 ms, α: 70°, matrix size: 64 × 64, 32 slices, voxel size 3×3×4 mm3). High resolution T1-weighted images were collected with a multiecho magnetization prepared gradient echo (MPRAGE) sequence, TR=2,300ms, TE=2.74ms, flip angle = 8 deg, matrix = 256×256×176 mm, voxel size = 1×1×1mm.
**2016 sample**^37^: MRI images in this sample were collected using a 3T Philips scanner, over two runs for a total scan time of 7 minutes 54 seconds. T2*-weighted images were collected using a gradient echo, echo-planar sequence (TR: 2,000 ms, TE: 29 ms, flip angle: 75deg, matrix size: 64 x 64 x 39, voxel size: 3.44 x 3.44 x 3.5mm^3^). High resolution T1-weighted images were collected with an MPRAGE sequence with the following parameters: TR/TE = 29/2,000 ms, flip angle=12 deg, matrix=256×256×160 mm, voxel size =1×1×1mm.

#### Task design

For the Filbey 2009 dataset^36^, the task consists of two runs of a pseudo randomized order of 12 tactile/visual stimulus presentations. Two types of stimuli are presented: (1) a marijuana cue (pipe, bong, blunt, joint), and (2) a neutral cue (pencil). Cues are presented for 20 s, followed by a 5 s rating period, during which craving ratings are self-reported on an 11-point scale. This is followed by a 20 s fixation period. The full task consists of a total of 12 pseudorandomized cue presentations. The task structure for the Filbey 2016 dataset^37^ is largely similar, but also includes a naturalistic cue (participant’s chosen fruit) for a total of 3 cue types and 18 presentations per run. Craving ratings are measured just as described above.

All 323 subjects (195 subjects with clinical label of chronic use) had two runs of data. Run length varied by the dataset from which the subject was taken. The subjects from the 2009 dataset had 281 TRs, and the subjects from the 2016 dataset had 405 TRs. For every subject, these TRs represented the totality of the run, including cue stimulus presentation periods, rating periods, and inter-trial intervals.

### fMRI data preparation

#### Preprocessing of fMRI

Results included in this manuscript come from preprocessing performed using FMRIPREP version stable^62^, a Nipype^63^ based tool. Each T1w (T1-weighted) volume was corrected for INU (intensity non-uniformity) using N4BiasFieldCorrection v2.1.0^64^ and skull-stripped using antsBrainExtraction.sh v2.1.0 (using the OASIS template). Brain surfaces were reconstructed using recon-all from FreeSurfer v6.0.1^65^, and the brain mask estimated previously was refined with a custom variation of the method to reconcile ANTs-derived and FreeSurfer-derived segmentations of the cortical gray-matter of Mindboggle^66^. Spatial normalization to the ICBM 152 Nonlinear Asymmetrical template version 2009c^67^ was performed through nonlinear registration with the antsRegistration tool of ANTs v2.1.0^68^, using brain-extracted versions of both T1w volume and template. Brain tissue segmentation of cerebrospinal fluid (CSF), white-matter (WM) and gray-matter (GM) was performed on the brain-extracted T1w using fast^69^ (FSL v5.0.9). Functional data was slice time corrected using 3dTshift from AFNI v16.2.07^70^ and motion corrected using mcflirt^71^ (FSL v5.0.9). This was followed by coregistration to the corresponding T1w using boundary-based registration^72^ with six degrees of freedom, using bbregister (FreeSurfer v6.0.1). Motion correcting transformations, BOLD-to-T1w transformation and T1w-to-template (MNI) warp were concatenated and applied in a single step using antsApplyTransforms (ANTs v2.1.0) using Lanczos interpolation. Physiological noise regressors were extracted applying CompCor^73^. Principal components were estimated for the two CompCor variants: temporal (tCompCor) and anatomical (aCompCor). A mask to exclude signals with cortical origin was obtained by eroding the brain mask, ensuring it only contained subcortical structures. Six tCompCor components were then calculated including only the top 5% variable voxels within that subcortical mask. For aCompCor, six components were calculated within the intersection of the subcortical mask and the union of CSF and WM masks calculated in T1w space, after their projection to the native space of each functional run. Framewise displacement^74^ was calculated for each functional run using the implementation of Nipype. Combined task/nuisance regression was then performed on the minimally preprocessed data using SPM12 (Wellcome Trust Centre for Neuroimaging). The nuisance regressor set consisted of the six realignment parameters, aCompCor regressors, discrete cosine-basis regressors, and a framewise displacement regressor. The task regressor set included onsets for marijuana cue presentation, marijuana cue rating period, control cue presentation, control cue rating period, and washouts for each cue. In addition, the Filbey 2016 dataset included regressors for fruit cue presentation and fruit cue rating period.

#### Parcellation

The noise-regressed voxelwise data were then parcellated using the Stanford functional ROIs for volumetric regions and networks, a highly validated scheme that is widely used for ROI-based and connectivity-based analyses^75^. The mean time series of each parcellated region was then computed by averaging the fMRI signal at every time point across voxels. This procedure served a dual purpose: first, it increased signal-to-noise ratio for relevant brain regions compared to voxel-based analyses. Second, it reduced the dimensionality of the data for subsequent analyses. The Stanford ROI atlas contains 90 regions, so the parcellation results in a 90 x (# of time points) matrix of whole brain activity for each subject.

#### Functional connectivity

Each region’s preprocessed time series was then correlated (Pearson) to all other regions’ time series. Pearson correlation automatically standardizes each region’s mean time series, so it is insensitive to differences in activation magnitude (i.e., scale) between the regions. Instead, it gives estimates of the pairwise timeseries activation similarities.

The decision to use parcellated functional connectivities was to 1) reduce the data dimensionality and the number of features relative to the number of observations, which is important in model fitting; 2) test the ability of network information to predict clinical label; and 3) improve our ability to subsequently interpret the fitted models, by using network analysis approaches. Further, functional connectivity has shown promise in other predictive modeling studies^31^. This approach each yielded a 646×90×90 matrix. To eliminate redundancy, only the upper triangles of the symmetric correlation matrices were retained (diagonal is each region’s correlation with itself), leading to a final vector input size of (90^2^ - 90)/2 = 4,005 features.

### Linear Classifiers

To train and evaluate classifiers, the full dataset was then divided into training and testing sets, using an 80/20 split: the training set included 516 samples (0.80 * 646) and the testing set included 130 (0.20 * 646). The training set was used for the 10-fold cross validated classifier training, hyperparameter optimization, and the final model selection. The testing set was set aside until the very end to test the out-of-sample fit of the four best performing models. The training-testing split was constructed to balance the overall clinical label (MJ or HC) proportions and include both runs of any subject completely in either the training or testing set. This part of the analytical workflow is shown in **Fig. 1b** and **1c**.

Four linear classifiers, namely L1- and L2-regularized Logistic Regression (LR) and L1- and L2-regularized linear kernel Support Vector Machine (SVM)^76^, were used to predict the target clinical label, i.e., chronic MJ use or not from the functional connectivity data. These classifiers were implemented using the scikit-learn Python package^77,78^. Generally, to separate classes, these classifiers learn a linear decision boundary in the feature space, generally referred to as a hyperplane, that then can be used to make class label predictions for new, out-of-sample data. In other words, the prediction (i.e., clinical label) is made based on the learnt weighted linear combination of the input features.

In particular, LR learns the logistic function that best fits the observations; the resulting sigmoid function gives the probabilities that each observation is in either class, which were thresholded at 0.5 in our implementation to produce the binary class predictions. In contrast, SVM learns a classification hyperplane that separates the two classes by the largest margin. In this case, the distances of the observations (each subject’s brain-wide pairwise functional correlations) to the hyperplane were converted to probabilities using Platt’s method, as implemented in scikit-learn^77,78^.

L_1_ and L_2_ regularization were used with both LR and SVM to penalize different types of information in the resultant models. L_1_ (“Lasso”) regularization penalizes the magnitudes of feature weights, and in doing so, produces a “sparse” feature space, such that only the features (e.g., region-region correlations) most informative to successful prediction will have a non-zero weight. Thus, L_1_ regularization reduces the number of features included in the model, which can improve interpretability, as well as predictive performance in case of many noisy and/or irrelevant features^79^. In contrast, L_2_ (“Ridge”) regularization penalizes the squares of squares of feature weights and minimizes their values, reducing their variance while retaining all of the features. This can improve prediction performance in cases where all the features can contribute useful information to the model. Various regularization strengths (α=1e-10, 1e-7, 1e-4, 0.1, 1, 10, 100, 1000) were tested in all classifiers, with larger strengths reflecting stronger penalization.

The four classification algorithms (L_1_- and L_2_-regularized LR and SVM), in combination with the values of *α* specified above, were evaluated in a 10-fold cross-validation setup (**Fig. 1b**). Here, the training set was randomly split into ten equally sized subsets (folds), stratified by class label to ensure the proportion of class labels was the same as in the larger dataset. Next, a model was trained on nine of the folds, and used to make predictions on the remaining tenth fold. This process was then repeated with each of the ten folds as the prediction set.

Each algorithm’s performance was then calculated by comparing the full prediction set with the available true labels of chronic MJ use in the training set. This performance was measured mainly in terms of prediction accuracy and the Area Under the Receiver Operating Characteristic (ROC) Curve (AUC) score^80^. ^80^In general, low regularization was found to have the highest cross-validated training performance. The α parameter resulting in the highest overall accuracy in the training set was selected for the subsequent analyses (α =1e-4). Finally, a model was learnt from the whole training set for each of the four classification algorithms. These models were then evaluated on the independent test set created earlier in terms of the evaluation measures mentioned above (**Fig. 1c**). Prediction accuracy, AUC, and weighted precision and recall measures^80^ were used for out-of-sample performance evaluation.

Each of the above final linear models returned a set of trained model weights that, when considered along with the values of the input features, are interpretable as the importances in determining the class label. These model weights were further explored using network analyses, as described below.

### Comparing functional activity to connectivity

To test our hypothesis that functional brain region correlations are more informative than more standard measures of activation magnitude, we also ran classification models with pairwise mean distances between activations of brain regions as model inputs. Three different mean distance controls were performed. In the first, the time series were averaged for each region and pairwise distances were calculated and used as the model inputs. In the second, the pairwise distances between regions’ average marijuana cue betas (from the task regression) were used. In the third, contrasts were generated between marijuana cues and control cues (marijuana > control), and the pairwise distances between regions’ contrast values were used. These control models were developed using the same process as described above for the functional connectivity inputs (features).

### Predictive importance analysis

Our next goal was to infer which functional correlations were most important to the predictions. Since linear models use the weighted sum of the weights and inputs to produce predictions, the contribution of a given functional correlation to the model’s predictions is given by the product of the model weight and the correlation values (i.e., the model weighted correlation values; see **Fig. 3**). For example, a positive weight and positive input (weight_pos_* correlation_pos_) makes a “positive” class prediction more likely, whereas a negative weight and positive input value makes a “negative” class (weight_neg_ * correlation_pos_) more likely, and so on. Similarly, the magnitude of a functional correlation’s contribution to classification depends on the magnitude of the product of the weight and input: a larger absolute value means a larger contribution of the functional correlation to the prediction.

Predictive importance analysis started by averaging the 90×90 connectivity matrices for each of the subjects in the training set used for classification model development. Next, the model weights were obtained for each linear model, and the element-wise product (i.e., Hadamard product) was computed between each model’s weight matrix and the group-averaged connectivity matrix to generate the weighted connectivity matrix expected to indicate the predictive importance of each pairwise connectivity.

For each row in this matrix, corresponding to the weighted connectivity vector associated with a particular region, the mean of the absolute values in the vector was calculated to represent the overall importance of the region’s weighted connectivity for the prediction of the clinical label. Four such scores for as many linear classification models were generated for each of the 90 regions and ranked by their average importance across all four models. Model-specific rankings were statistically compared using Kendall’s tau to assess correspondence between each pair of models. The top twenty regions of highest weighted connectivity, or highest predictive importance, were selected to visually examine their individual connectivity patterns and corresponding weights.

Region-specific predictive importance scores were validated by comparing them to meta-analytic uniformity and association maps retrieved from Neurosynth^46^. The keyword ‘craving’ was used to yield maps aggregated from 80 published studies. The uniformity and association maps each provide unique information; the uniformity map displays regions of consistent activation across all studies, while the association map displays regions that are active over and above maps from other keywords. Maps were thresholded at p<0.01 with FDR correction and projected on the Stanford functional ROIs. For each ROI, the proportion of non-zero voxels, and average non-zero signal was calculated, indicating average ROI activity in the meta-analytic map. Finally, these scores were thresholded to limit reporting of voxel activity in regions that contained too few active voxels. Given the relative sparsity of the association map compared to the uniformity map, the former was thresholded at 5% voxel participation and the latter at 25%.

### Network Analysis

Given the fact that we used functional connectivity as our input features, we then used network analysis to analyze the distributed patterns of the connectivity important for prediction.

The first step in preparing weighted connectivity data for network analysis was to threshold subject-specific matrices. Thresholding is a commonly used strategy in network neuroscience to remove spurious network connections, and improve stability and modularity of network features^81–84^. The absolute values of the weighted connectivity values were taken, as the magnitudes of weighted connectivity set as the strength of node-to-node connections in the graph. The transformed matrix was used to generate a sparse graph, where nodes represented regions, and the edges represented the strength (i.e., importance) of connectivity values between two regions to prediction.

Subject-specific weighted connectivity matrices were treated as adjacency matrices corresponding to an undirected weighted graph. We binarized the dense weighted connectivity matrix at 2% density (top 2% of values converted to 1 and others to 0) to improve signal-to-noise ratio and remove weak connectivity strengths. The resultant binary matrix was used to calculate the node- and graph-level properties. Finally, we generate the graph structure by considering the binarized weighted connectivity matrix as an adjacency matrix using the networkx package in Python^85^.

With a unique graph structure for each subject, we calculate subject-specific degree centrality (DC), a node-level graph property (see **Fig. 6a**), which refers to the fraction of nodes to which a particular node is connected, normalized by dividing by the maximum possible connections. In this graph, each node represents a brain region and connection edges between two nodes represent the importance of the connectivity between those two nodes for the classifier. Thus, nodes with high degree centrality can be considered to be brain regions whose connectivities to other regions help the classifier distinguish chronic MJ users from non-users. Conversely, graph isolates are defined as nodes with lowest degree centrality across participants. In other words, they are brain regions whose connectivities to other regions do not help the classifier distinguish between chronic users and healthy controls. DC calculation was performed using networkx’s degree_centrality function, which accepts a graph structure and calculates the DC of each node. For each brain region (i.e., node in the network), the distribution of DC for that region was calculated across all participants. Regional DC scores were ordered by highest median score.

Next, graph-level metrics of the connectivity matrices were calculated next by deriving global efficiency scores at a subject-specific level (see **Fig. 6c**). The efficiency between two nodes is defined as the inverse of the shortest path between them, providing an overall measure of the ability of a network to propagate information effectively. Efficiency metrics were calculated using the built-in networkx function ‘global_efficiency’. Two-sample Mann-Whitney U tests were performed to test for differences in median efficiency scores between users and non-users.

Finally, communities with high predictive importance in classifying chronic marijuana use were identified (see **Fig. 6b**). First, a group-average weighted correlation matrix was calculated by taking the mean of all un-thresholded subject-specific weighted connectivity matrices calculated above. This mean correlation matrix was then thresholded at 2% density, generating the group-average graph structure. Then, the Girvan-Newman hierarchical community-detection algorithm was used to detect clusters that were highly interconnected within the graph. Briefly, the Girvan-Newman algorithm iterates between the following steps: (1) edge betweenness, defined as number of paths between all nodes that include a particular edge, is calculated for each edge; (2) edges with the highest betweenness are removed; (3) betweenness of all the edges is recalculated. The final communities are defined as the node clusters comprised of nodes that are highly connected within the cluster, but sparsely connected to other clusters. The nodes (rows) in the original weighted correlation matrix were then reordered based on identified community structure to reveal modular clusters. The communities were then ranked by their average degree centrality score, with those having the highest ranks defined as the most predictively important.

The predictive importance of the communities was corroborated using the following stepwise prediction approach.

Starting with the highest ranked community, we tested the classification accuracy of the trained model using correlations obtained from that community. In each subsequent step, the correlations from the next highest ranked community were added. For each subject, the non-redundant correlations of the regions within that community to all other regions were used to generate the distances to the decision hyperplane. These distances were generated by taking the dot product of those correlations and their trained model weights and adding the intercept from the whole trained model. With each additional community, we selected the optimal decision threshold for that iteration as the one with the highest prediction accuracy in the training data. This threshold was then applied to the testing data and prediction accuracy was reported (see **Fig. 9**). The best performing subset of communities was determined by the testing set accuracy. To determine whether these accuracies were a function of the unique, included communities or just the number of pairwise functional correlations, a permutation approach was used. 1000 permutations were computed using the same approach as described above, except randomly shuffling the regions included in each community while preserving the number of pairwise correlations included at each step. The permutation p-value was calculated as the percentile of the best performing non-permuted accuracy in the distribution of the 1000 permuted accuracies at that same step and was also reported in **Fig. 9**.

## DATA AND CODE AVAILABILITY

All the code related to analyses in this study is publicly available at https://github.com/kulkarnik/mj_classifier. The data are available in the same repository.

## AUTHOR CONTRIBUTIONS

K.K. and M.S. conceptualized and designed the predictive-explanatory modeling framework, carried out the implementation, and analyzed the data. G.P. provided feedback on modeling framework. V.C., F.F., K.H., and G.P. contributed to the interpretation of the results. K.K. and M.S. wrote the manuscript with critical feedback from all authors, especially G.P. F.F. and K.H. collected and organized the data. X.G., D.S., and G.P. supervised the project.

## ACKNOWLEDGEMENTS

The authors acknowledge support by the US National Institutes on Drug Abuse under awards R01 DA043695 and R21 DA0492243. The authors also acknowledge the computational resources and staff expertise provided by Scientific Computing at the Icahn School of Medicine at Mount Sinai. The content is solely the responsibility of the authors and does not necessarily represent the official views of the National Institute on Drug Abuse.

